# Simultaneously vocalizing Asian barbets adopt different frequencies without coordinating temporal rhythms

**DOI:** 10.1101/754580

**Authors:** Anand Krishnan

## Abstract

Sound stream segregation is an important challenge faced by simultaneously vocalizing animals. Chorusing birds, for instance, coordinate vocal timing to minimize overlap. Alternatively, other birds may use frequency differences to segregate sound streams, and vocalizing at different frequencies may enable them to remain distinct from each other. Here, I show that conspecific Asian barbets vocalize at distinctly different peak frequencies from each other. Additionally, they also differ in repetition rate, as measured by the inter-phrase interval. However, conspecific individuals across species do not temporally coordinate with each other during vocal interactions, maintaining independent and highly stereotyped individual rhythms together with different peak frequencies. Frequency differences between individuals may facilitate sound stream segregation when calls overlap in time. I propose that frequency differences between conspecifics may be widespread among birds possessing stereotyped, repetitive calls such as those found in barbets. This may enable segregation of competing sound streams both during cooperative duets and competitive singing during territorial interactions.

## Introduction

Animal choruses represent a striking natural example of the ‘cocktail party problem’, where individuals must segregate relevant information from competing streams of sound, both conspecific and heterospecific (Bee and Micheyl, 2008; Greenfield, 1994). This applies both to territorially singing males, and birds singing agonistically together in duets (Hall, 2004; Hall, 2009). In both scenarios, conspecific individuals must emit distinct signals in order to communicate with each other. Vocalizing at differing frequencies (Krishnan and Tamma, 2016; Narins, 1995; Nityananda and Bee, 2011), or altering vocal timing (Brumm, 2006; Cody and Brown, 1969; Fleischer et al., 1985; Luther, 2008) may minimize masking interference from overlapping sounds. Some birds are known to simply sing together without temporal coordination, and their vocalizations drift in and out of phase with each other (Hall, 2009; Payne and Skinner, 1970). As a result, simultaneously vocalizing conspecifics may repeatedly overlap in time, depending on the differences in repetition rate. In such cases, frequency differences may assume greater importance in sound stream segregation (MacDougall-Shackleton et al., 1998). By recording animal choruses and isolating the frequencies and repetition rates of each individual, we may obtain an indication into how each individual may be able to distinguish themselves over a chorus of conspecifics.

Here, I describe the vocal strategies employed by Asian barbet (Piciformes: Megalaimidae) (Short and Horne, 2001) choruses in India. Across Asia, multiple species of sympatric barbet co-occur, each vocalizing in a species-specific frequency band (Krishnan and Tamma, 2016). In addition to these interspecific differences, multiple individuals of the same species frequently vocalize together in choruses (Ali and Ripley, 1997; Short and Horne, 2001). However, the mechanisms that barbets employ to segregate their signals from conspecifics remain poorly understood. By studying simultaneously vocalizing conspecifics of four different species, I aimed to preliminarily understand whether vocal patterns were similar or diverse across the family. My study examined whether simultaneously vocalizing conspecific barbets of each species segregate from each other either in vocal peak frequencies, or by coordination of vocal timing. For this, I examined whether conspecifics vocalizing together differed in peak frequency and repetition rates, and then examined the inter-phrase intervals between two conspecific individuals to elucidate whether, across species, there was any evidence that barbets coordinated their calls with respect to vocal conspecifics. Understanding the vocal strategies employed by these non-passerine birds enables a broader understanding of the diversity of animal strategies to minimize acoustic masking.

## Materials and Methods

### Recording

I recorded barbet choruses in the city of Pune in Maharashtra (Peninsular India), and the village of Mandal in Uttarakhand (Western Himalayas) in March-April 2018, early in the breeding season. Each site houses two species of barbet, *Psilopogon viridis/P.haemacephalus* in Pune, and *P.virens/P.asiaticus* in Mandal (Figure 1A. For recordings, I used Sennheiser (Wedemark, Germany) ME62 omnidirectional microphones connected to a Zoom H6 (Tokyo, Japan) recorder sampling at 44.1KHz, making note of multiple simultaneously vocalizing conspecific individuals. The recorder and microphones were stationary on the ground to avoid movement noise. The overall dataset consisted of approximately seven hours of barbet chorus recordings.

**Figure 1:**
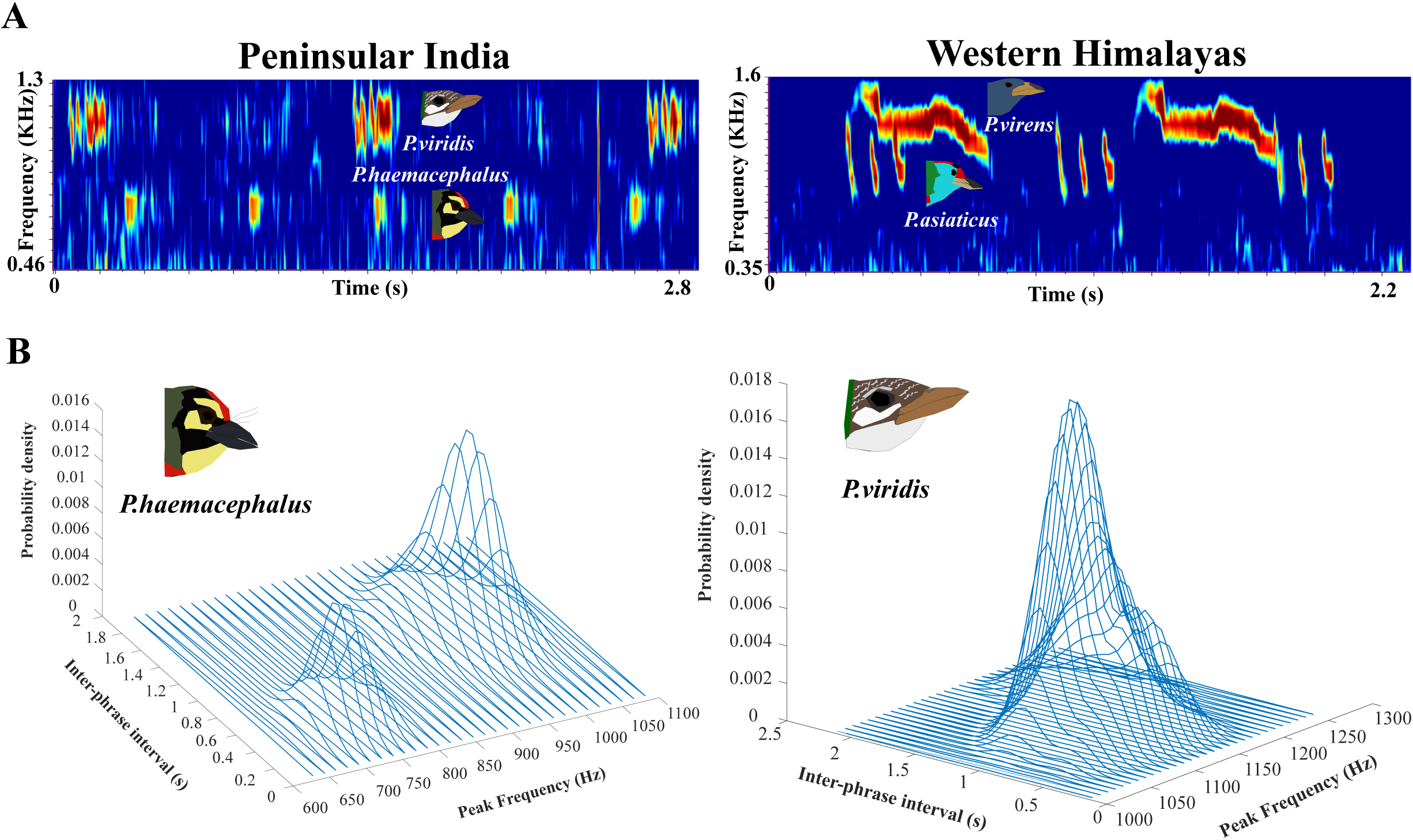
**A.** Spectrograms of barbet vocalizations from Peninsular India (left) and the Western Himalayas (right). **B.** Three-dimensional probability density distributions indicating the occurrence of different mean PF and repetition rates (IPI) for *P.haemacephalus* and *P.viridis*.

### Analysis

Using Raven Pro 1.5 (Cornell Laboratory of Ornithology, Ithaca, NY, USA), I digitized vocalizations of barbets from each recording and calculated peak frequencies of each note (PF, using an FFT window size of 2048 for higher frequency resolution of small differences) as well as the inter-phrase interval (IPI, the time gap between the end of one phrase and the beginning of the next, using a window size of 512 for higher resolution of small timing differences). For each species, I digitized approximately 2500 individual phrases (1-5 notes/phrase depending on the species). I listened to all recordings using headphones while labeling vocalizations, differentiating conspecific individuals by the differences in relative amplitude (birds that were further from the microphone were lower-amplitude in recordings, and distinguishable in Raven Pro). First, I examined whether two simultaneously vocalizing conspecific individuals differed in PF and IPI, using paired Wilcoxon signed-rank tests. Next, I measured the interquartile range or IQR of IPI (for 108 total instances of a vocal barbet across all four species), to quantify stereotypy in temporal rhythms. Secondly, to examine whether simultaneously vocalizing conspecifics (36 instances across four species, an instance defined as bouts of vocalizations from the same individual without long gaps; this was the most conservative way to identify behavioral instances given that I could not identify individual birds in the field) were temporally synchronous or asynchronous with each other, I calculated the time lag between the beginning of each phrase of one individual and the closest call of the other using MATLAB (Mathworks Inc., Natick, MA, USA). If birds were synchronous, I predicted that the distribution of time lags across a bout should show a clear peak and low coefficient of variation (i.e. a stereotypical time lag between individuals) (Taylor et al., 2019). However, an asynchronous bout would imply independent rhythms from each other; the two individuals would thus drift in and out of phase, resulting in a uniform distribution of time lags. For each instance with 5 or more measured time lags (27 in total), I performed two-sample Kolmogorov-Smirnov tests against 100 randomly generated uniform distributions spanning the same range of values. I scored the results as 0 if they did not differ significantly from uniform (P>0.05), or 1 if they did (P<0.05), and measured the percentage of total 1’s (out of 100 tests) for each of the 27 instances. A score of 0 for an instance indicated a uniform distribution of time lags consistent with independent temporal rhythms, and a score of 1 indicated a constant time lag consistent with two birds singing in coordination.

## Results

### Conspecific barbets sing at different frequencies and repetition rates

PF-IPI probability density plots revealed intraspecific variation in PF and IPI (Figure 1B, see supplementary videos for 3D rotations). For example, in *P. haemacephalus*, two distinct peaks were consistent with two distinct types of song, one at about 0.7KHz with an IPI at approximately 0.5s, and a slower song at approximately 0.95KHz with an IPI over 1s (total of 27 measured instances). For *P. viridis*, I observed a similar bimodal distribution with variation (difference between minimum and maximum) of over 130Hz in PF and over 1.3s in IPI (total of 65 measured instances). For *P.virens* and *P.asiaticus*, my data likely involved a long series of measurements from relatively few individuals. Thus, although I digitized a similar number of phrases in these species, these were separable into only 9 and 7 separate behavioral instances, respectively. Although this sample size was too small to construct probability density distributions, the measured PF and IPI exhibited high intraspecific variation (maximum-minimum) for both species: 116Hz PF and 0.23s IPI variation for *P.virens*, and 148Hz PF and 0.04s IPI variation for *P.asiaticus* (although IPI variation was small in the latter species, there was a consistent difference in the few cases when two individuals were vocalizing together, even across very long bouts, as discussed in the next section). Intraspecific PF and/or IPI variation was thus broadly consistent across all four species, suggesting that simultaneously vocal conspecifics may adopt different frequencies and/or repetition rates from each other.

To statistically test this assertion, I compared the PFs and IPIs of simultaneously vocal barbets using paired Wilcoxon signed-rank tests. Because the data for *P.virens* and *P.asiaticus* consisted mainly of very long interactions across relatively few individuals (therefore, a very large number of labeled notes, but few individual instances of simultaneously vocal barbets), I performed this analysis only for *P. viridis* (N=22 instances of two barbets vocalizing together) and *P.haemacephalus* (N=8). In both species, tests revealed significant (*P. viridis*: IPI: signed rank= 253, P<0.001, PF: signed rank= 256, P<0.001, *P.haemacephalus*: IPI: signed rank= 36, P<0.01, PF: signed rank= 36, P<0.01). This suggests that intraspecific variation in the measured time and frequency parameters resulted from conspecific individuals adopting different frequencies and vocal rhythms from each other. This can be seen in Figure 2A, where the spectrograms clearly show conspecifics vocalizing at different frequencies and repetition rates.

**Figure 2:**
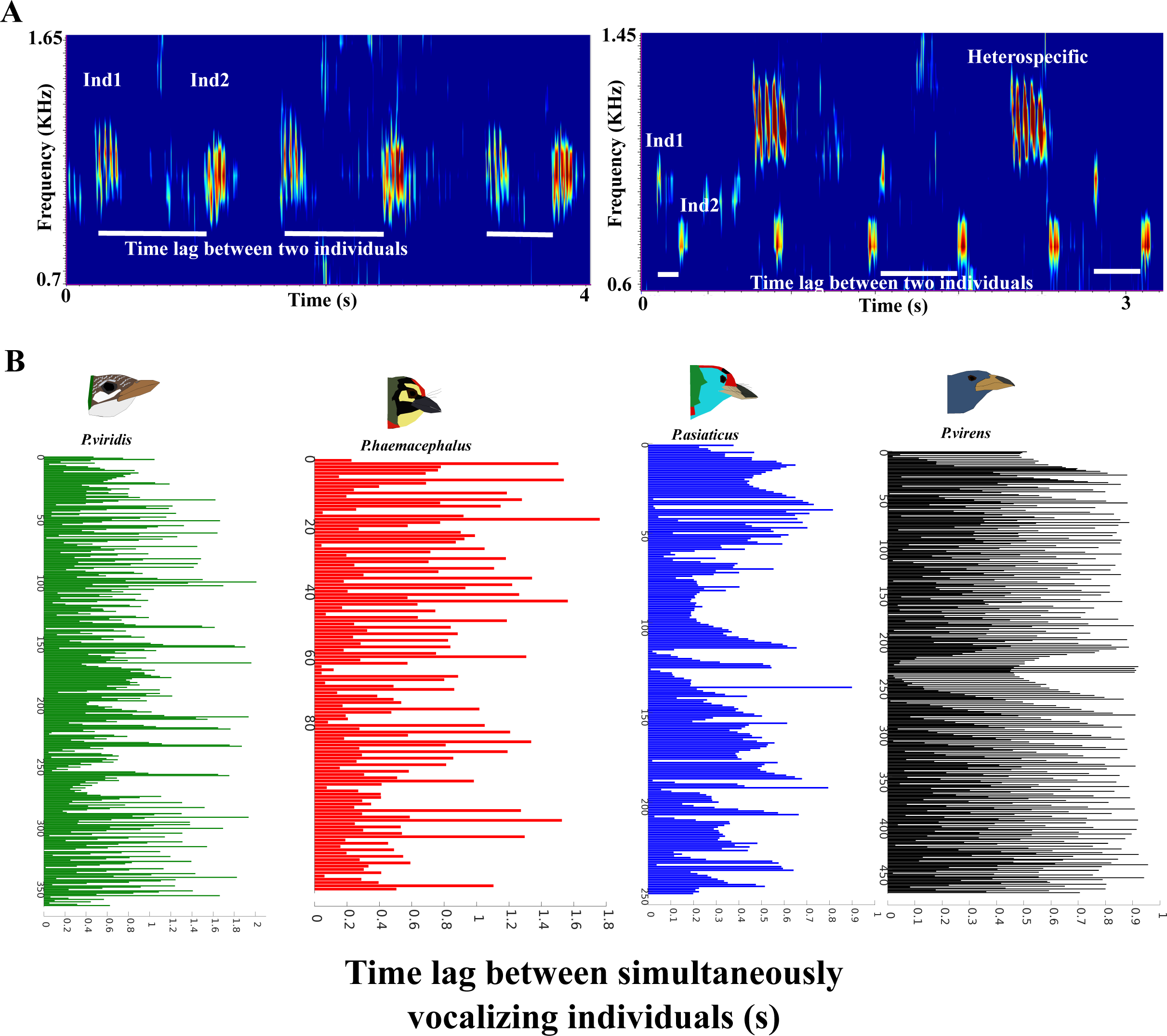
**A.** Spectrograms of two simultaneously vocalizing *P.viridis* (left) and *P.haemacephalus* (right), demonstrating intraspecific differences in frequency and repetition rate. The white bars represent the time lag between two individuals; note how it changes with each repetition. **B.** Graphs of all measured values of time lag between two individuals for each species of barbet. The longer the horizontal bar, the longer the time lag; the y-axis represents the number of such measurements made per species. Values vary from approximately 100ms to over a second, indicating a lack of temporal synchrony between individuals.

### Simultaneously vocalizing conspecifics sing at independent rhythms without synchronization

I next sought to establish whether vocally interacting conspecific barbets adjusted their repetition rates to maintain synchrony with their vocal neighbors. Across 36 instances of simultaneously vocalizing conspecifics, I calculated the time lags between the calls of two individuals (Figure 2A). For this, I used data from all four species, as there were enough individual vocalizations digitized for the sample size of total time lag values to be sufficient for statistical analysis. The time lags between the calls of two individuals had very high coefficients of variation for each species (*P.haemacephalus:* 144.13%, *P.viridis:* 164.52%, *P.virens:* 182.55%, *P.asiaticus:* 197.11%), supporting a lack of vocal timing coordination (Hall, 2009). Time lags exhibited a range of values across all 4 species, as opposed to a single value that would be expected if two individuals coordinated vocal timing by phase-locking (Figure 2B). When compared to 100 randomized uniform distributions (this was done for each of 27 instances, see Methods), time lags fit a uniform distribution 92% of the time on average (average score 0.08, P>0.05, Supplementary Data, Figure 3A, also see Methods). This uniform distribution of time lags, together with high CVs, thus supports barbets vocalizing with independent temporal rhythms rather than synchronizing with each other. In further support of this, each barbet maintained its individual rhythm, whether vocalizing alone or with conspecifics.

**Figure 3:**
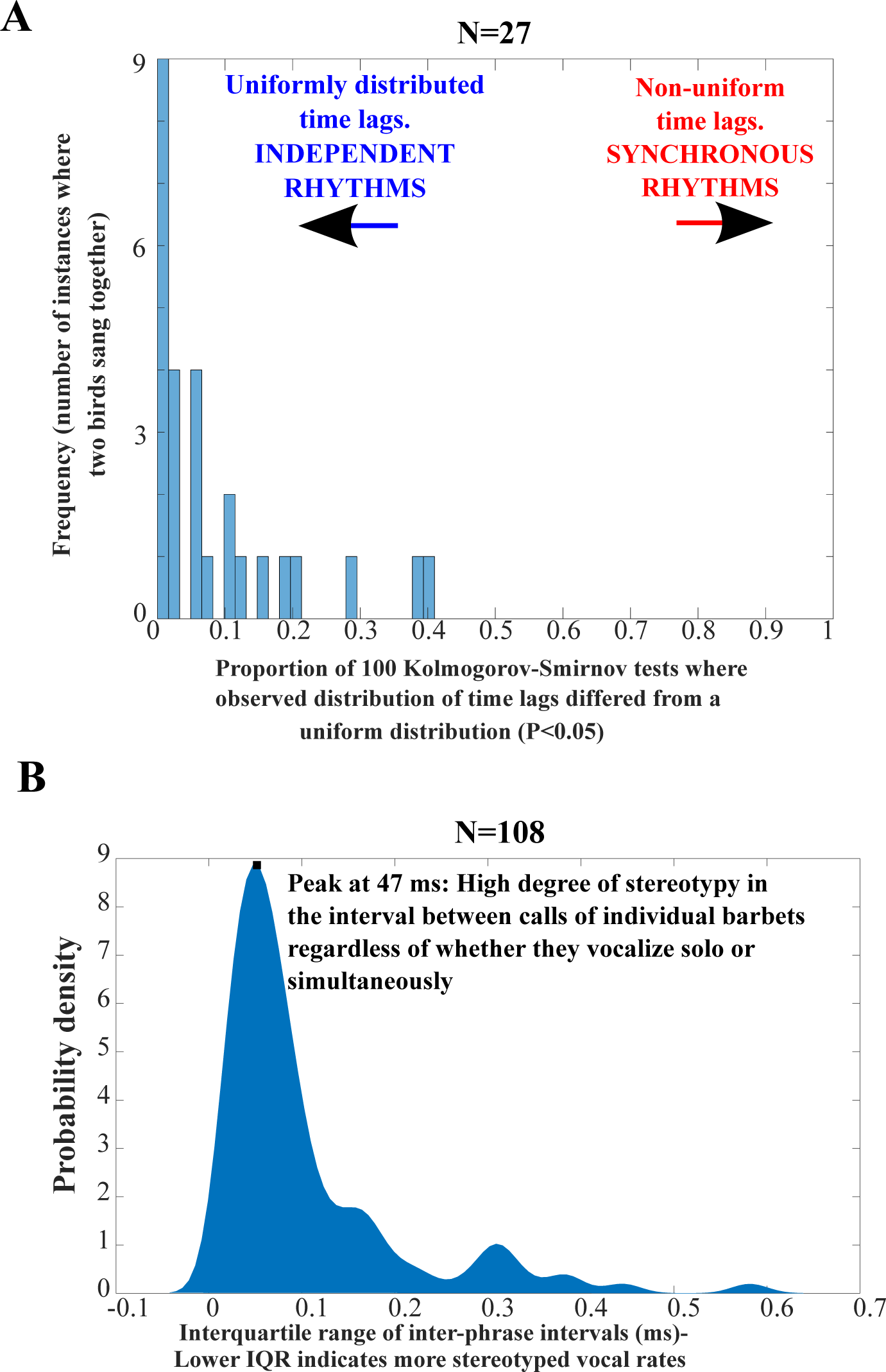
**A.** Proportion of 100 Kolmogorov-Smirnov tests in which the observed distribution of time lags differed from a random uniform distribution (for 27 instances of two conspecific barbets vocalizing together). In 18/27 cases, the observed distribution fit >90% to a uniform distribution (x-axis values <0.1, indicating independent temporal rhythms without synchrony). Almost all other instances also fit well to a uniform distribution (the lowest being a 59% fit to uniform, x-axis value 0.41), again supporting independent vocal rhythms. **B.** Distribution of IQRs for the IPI of each individual instance of a vocalizing barbet across the entire dataset (both solo and simultaneous calling, N=108). The peak at 47ms indicates very low IQRs, and therefore a highly stereotyped temporal rhythm regardless of whether the barbet is vocalizing solo or with conspecifics.

The probability density plot of IQRs for the IPI of each individual barbet (all individual behavioral instances measured, either solo or with another barbet) peaked at 47ms (Figure 3B), indicating <50ms jitter in vocal timing across all species (including measurement-related variation). This analysis included every individual instance of a vocal barbet across the dataset (solo and with other conspecifics), in spite of which I observed a sharp peak indicating a precise vocal rhythm. This further supports the idea that barbets maintain stable, independent vocal rhythms regardless of whether they vocalize solo or with conspecifics.

## Discussion

Avian vocal interactions have received much study for the precisely coordinated vocal timing between individuals; many of these studies have focused on duetting birds (Fortune et al., 2011; Hall, 2009), but others have also focused on temporal asynchrony of songs to minimize overlap (Cody and Brown, 1969; Fleischer et al., 1985). However, other birds exhibit independent temporal rhythms with no phase-locking or coordination between simultaneously vocalizing individuals (Hall, 2009). My data suggests that Asian barbets do not coordinate vocal timing with respect to other vocal conspecifics, and I also find that simultaneously vocalizing individuals tend to differ in the peak frequencies of their vocalizations. If two barbets vocalize at independent and different rhythms, their vocalizations will drift in and out of phase with each other (resulting in a uniform distribution of time lags between the two individuals, as opposed to the single peak one might expect if they were coordinated with each other).

Dilger (Dilger, 1953) described simultaneous singing in *P.haemacephalus* involving both members of a pair vocalizing at different frequencies. My dataset, consisting of broad passive recordings of barbet choruses, presumably consists of a mix of vocal individuals, both vocal pairs and territorial countersinging birds. This, of course, assumes that both members of a pair sing across species as per Dilger’s observations, which requires further study. Because barbets are often difficult to locate when singing, usually high up in the canopy, and the sexes are alike, it is extremely challenging to identify individual birds. However, at the species-level, I find that conspecific individuals vocalize together with differences both in peak frequency and the interval between phrases. Additionally, in no instance do I find evidence of birds changing their temporal rhythms to minimize overlap. Instead, barbets appear to simply adopt different temporal rhythms from each other, which may involve paying attention only to the start of another bird’s bout. Although different repetition rates may reduce temporal overlap to some extent, some vocalizations of two individuals will still overlap in time. In this case, frequency differences between individuals may support sound stream segregation (MacDougall-Shackleton et al., 1998; Nityananda and Bee, 2011). Frequency differences between conspecifics are smaller than those between heterospecifics (Krishnan and Tamma, 2016), suggesting that they serve to differentiate individuals within each species’ frequency band.

Some species of the related African barbets (Lybiidae:*Trachyphonus*) pair-duet, and certain species appear to exhibit vocal timing coordination (Horne and Short, 1983; Thorpe, 1963). However, duetting partners in other species may exhibit independent rhythms (Payne and Skinner, 1970). It is thus possible that Asian barbets may sometimes coordinate their rhythms over short time scales, although my study does not uncover evidence of this. By and large, the evidence I present suggests that frequency differences between conspecifics are more important than temporal mechanisms to minimize vocal overlap. Similar mechanisms may also operate in other non-passerine birds such as the pheasant-coucal, where simultaneously vocalizing members of a pair exhibit different frequencies (Maurer et al., 2008). Comparative study may thus help us understand the diversity of strategies employed by birds to minimize masking interference when singing in a chorus.

## Acknowledgments

I thank Rohit Chakravarty, Zareef Khan, Baseer Baniya and Arun Varghese for assistance during recordings, Raghav Rajan and his lab for inputs on the data, Samira Agnihotri and Shivam Chitnis for discussions on analysis.

## Funding

My research is funded by an INSPIRE Faculty Award from the Department of Science and Technology, Government of India and an Early Career Research (ECR/2017/001527) Grant from the Science and Engineering Research Board (SERB), Government of India.

